# Microfluidic *Streptomyces* Cultivation for Whole Lifecycle Characterization and Phenotypic Assays Enabled by Nanogap-stabilized Air-Water Interface

**DOI:** 10.1101/2021.05.13.443940

**Authors:** Dongwei Chen, Mengyue Nie, Wei Tang, Yuwei Zhang, Yuxin Qiao, Juanli Yun, Jian Wang, Ying Lan, Liang Ma, Yihua Chen, Wenbin Du

**Author notes:** **Address correspondence to** Liang Ma,; Yihua Chen, or Wenbin Du,.

## Abstract

*Streptomyces* is a model filamentous prokaryote to study multicellular differentiation and a rich reservoir for antibiotics discovery. In their natural conditions, *Streptomyces* grows at the interface of porous soil, air, and water. The morphological development of *Streptomyces* is traditionally performed on agar plates and mostly studied at the population levels. However, the detailed lifecycle of *Streptomyces* has not been well studied due to its complexity and lack of research tools which can mimic their natural conditions in the soil. Here, we developed a simple assembled microfluidic device for cultivation and the entire lifecycle observation of *Streptomyces* development from single-cell level. The microfluidic device composed of a microchannel for loading samples and supplying nutrients, microwell arrays for seeding and growth of single spores, and air-filled chambers aside of the microwells that facilitate growth of aerial hyphae and spores. A unique feature of this device is that each microwell is surrounded by a 1.5 µm gap connected to an air-filled chamber which provide stabilized water-air interface. We used this device to observe the development of single *Streptomyces* spores and found that unlike those in bulk liquid culture, *Streptomyces* can differentiate at water-air interfaces in microscale liquid culture. Finally, we demonstrated that phenotypic A-Factor assay can be performed at defined time point of its lifecycle. This microfluidic device could become a robust tool for studying *Streptomyces* multi-cellular differentiation and interaction at single cell level.

**IMPORTANCE:** We describe a microfluidic device that mimics the natural porous environment for the growth and development of *Streptomyces*, the model system for bacterial multicellularity. The microfluidic device is used for cultivation and the entire lifecycle observation of *Streptomyces* development from single-cell level, including growth of aerial filaments. The aerial hyphae development of *Streptomyces* at the water-air interface was observed at real time in the microfluidic device. The early growth, opportunistic transformation (in the gap), and merging of aerial hyphae of *Streptomyces* in the microfluidic device were observed for the first time. It will play an important role in finding single-cell heterogeneity to study secondary metabolites related to the complex lifecycle of *Streptomyces*.

## INTRODUCTION

Streptomycetes are gram-positive filamentous bacteria that play crucial roles in their habitat because of their broad range of metabolic processes and biotransformation including degradation of chitin and cellulose (1-3). They are the most important natural source of bioactive compounds such as antibiotics and anti-tumor agents, producing two-thirds of the antibiotics of medical and agricultural interest (4-6). In nature, *Streptomyces* grows primarily in soil with porous structures that retain water in the micron-sized cavities and channels. Nutrients, oxygen, water transport and other environmental factors may have profound impact on the their physiology, morphological development and outset of secondary metabolism (7). Although recent research with advanced genomic tools has made great progress in uncovering their genetic potential, a lot of discovered pathways are cryptic, which means they are either silent or poorly expressed for those grown on agar plate or in liquid media in the standard laboratory conditions, presumably due to the inability to recreate the microscale porous structure as well as nutritional and environmental circumstances in their natural soil habitat. In addition, bulk cultivation method only allows spatial control down to millimeter scale, and is often not convenient for nutrient and chemical exchange.

Microfluidics has emerged as a new tool to study microbiology because it offers many advantages including micrometer-scale spatial resolution and flexible temporal control of nutrients exchange and chemical gradients (8). Microfluidic tools have been used to study microbiology in many ways such as single-cell isolation and manipulation, bacterial chemotaxis, quorum sensing, and population dynamics (9). Although high-throughput enrichment and sorting of soil-derived *Actinobacteria* in microfluidic droplets have been described (10), the development and differentiation of *Streptomyces* using microfluidic chips are rarely reported until 2016 (11). The challenge in *Streptomyces* cultivation is that their growth and differentiation rely on a stabilized water-air interface. When cultivated on a solid agar, *Streptomyces* have a typical lifecycle including germinate of vegetative hyphae in the solid substrate, forming of hydrophobic aerial hyphae, and development of airborne spores which allow dispersion (12). However, in a standard liquid medium, *Streptomyces* mainly exists as vegetative hyphae that tangle together to form many small pellets and clumps with very few aerial hyphae (13). Therefore, direct miniaturization of standard liquid culture in a microchamber without a stabilized water-air interface is not a generalizable method for cultivation of *Streptomyces*.

To overcome these challenges, we describe a microfluidic device integrating liquid containing microwells and air-filled chambers to establish a stabilized water-air interface for cultivation, the whole lifecycle observation of *Streptomyces* differentiation and phenotypic assay. The device can achieve micron-scale spatial resolution, maintain culture conditions over an extended period of time with optional nutrient and chemical exchange, and enable single-cell cultivation and observation; thus, it is a useful tool for exploring *Streptomyces*’s development and behavior under controlled circumstances. The microfluidic platform was validated by culturing two representative *Streptomyces* strains, and a phenotypic assay with A-Factor. A-Factor and analogues are autoregulatory factors involved in secondary metabolism and/or morphological differentiation in Actinomycetes (14). They are essential for the aerial hyphae formation in *Streptomyces* (Fig. S3A). However, previous studies showed that the timing of A-Factor introduction is important for morphogenesis and secondary metabolism (15). The nutrient exchange channel incorporated in the microfluidic chip allows the A-Factor assay to be performed at defined time points in the lifecycle with mutant organism that is deficient in A-Factor synthesis.

## RESULTS AND DISCUSSION

### Design of the microfluidic device

We designed a microfluidic device with an array of microwells for the entire lifecycle observation of *Streptomyces* including cultivation and observation. This device incorporates two important design features: i) stable water-air interface enabled by nanogaps; ii) well controlled nutrient and chemical exchange through a microchannel. *Streptomyces* undergoes complex lifecycles involving vegetative hyphae growth in substrates and aerial hyphae growth in the air. We mimicked the natural microenvironment by assembling liquid-containing microwells and air-filled chambers for vegetative growth and aerial growth, respectively. The gap between two assembled glass plates is 1.5 μm, which is slightly larger than the diameter of the hyphae (1 μm). This separation ensures that the aerial hyphae can readily pass through the gap (Fig. 1A). The chamber is 15 μm in height so that the aerial hyphae have sufficient space to keep a natural state. When the microwells are filled with liquid, water-air interfaces are formed between the microwells and the air-filled chambers. The glass plates offer a reliable barrier to minimize evaporation. The plates also exhibit hydrophobic surface after treatment with fluorosilane, and thus the surface tension was relatively high (Fig. 2B).

**FIG 1.**
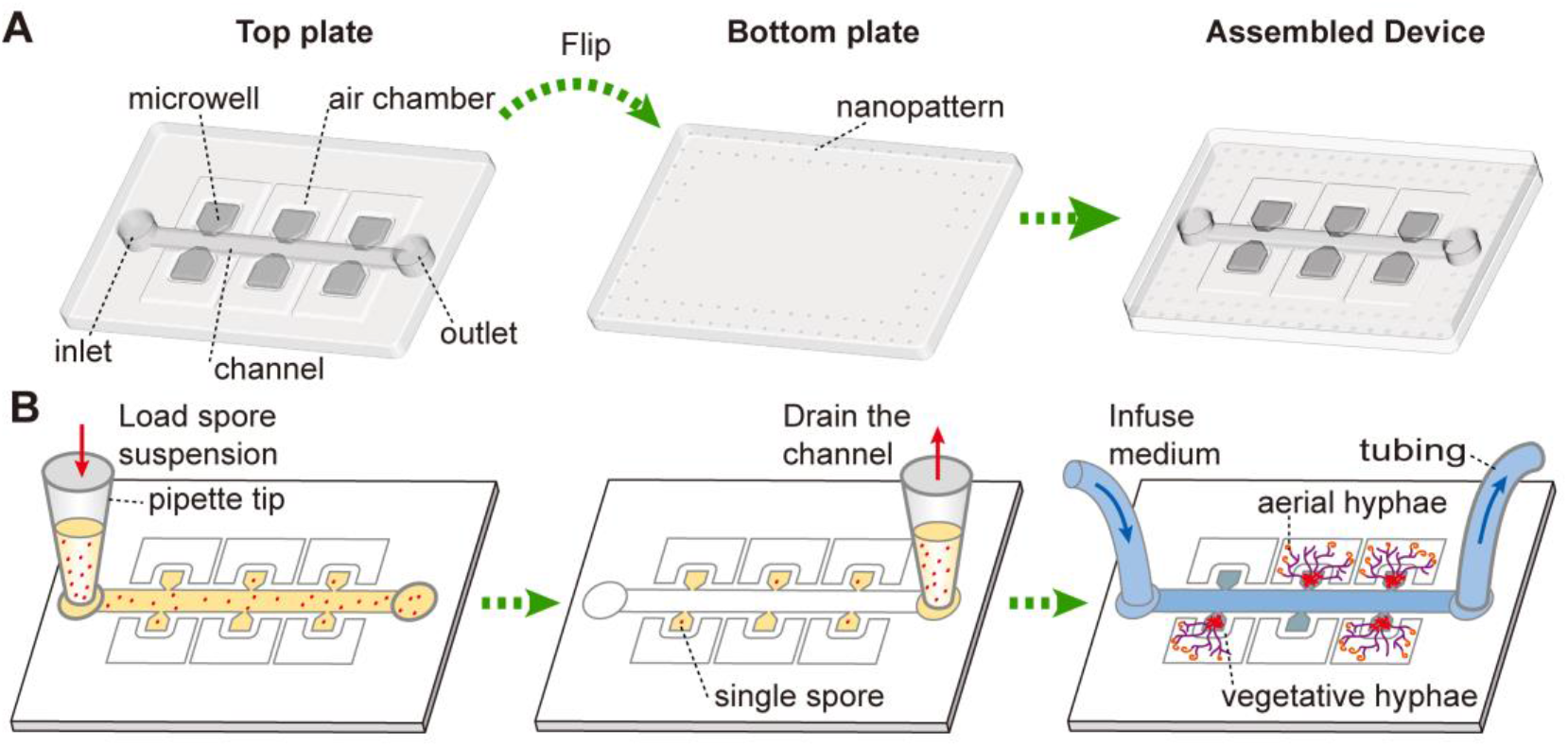
Illustration of the microfluidic chip for lifespan observation of *Streptomyces*. (A) Assembly and setup of the device. (B) Spore suspension is loaded into the microwells by a pipette. Concentration of spores is controlled to allow single spore trapping in the microwells based on a Poisson distribution. The channel was drained by pipette to remove spores in the channel. Culture medium was continuously infused into the device. The *Streptomyces* development process was observed by an inverted microscope.

**FIG 2.**
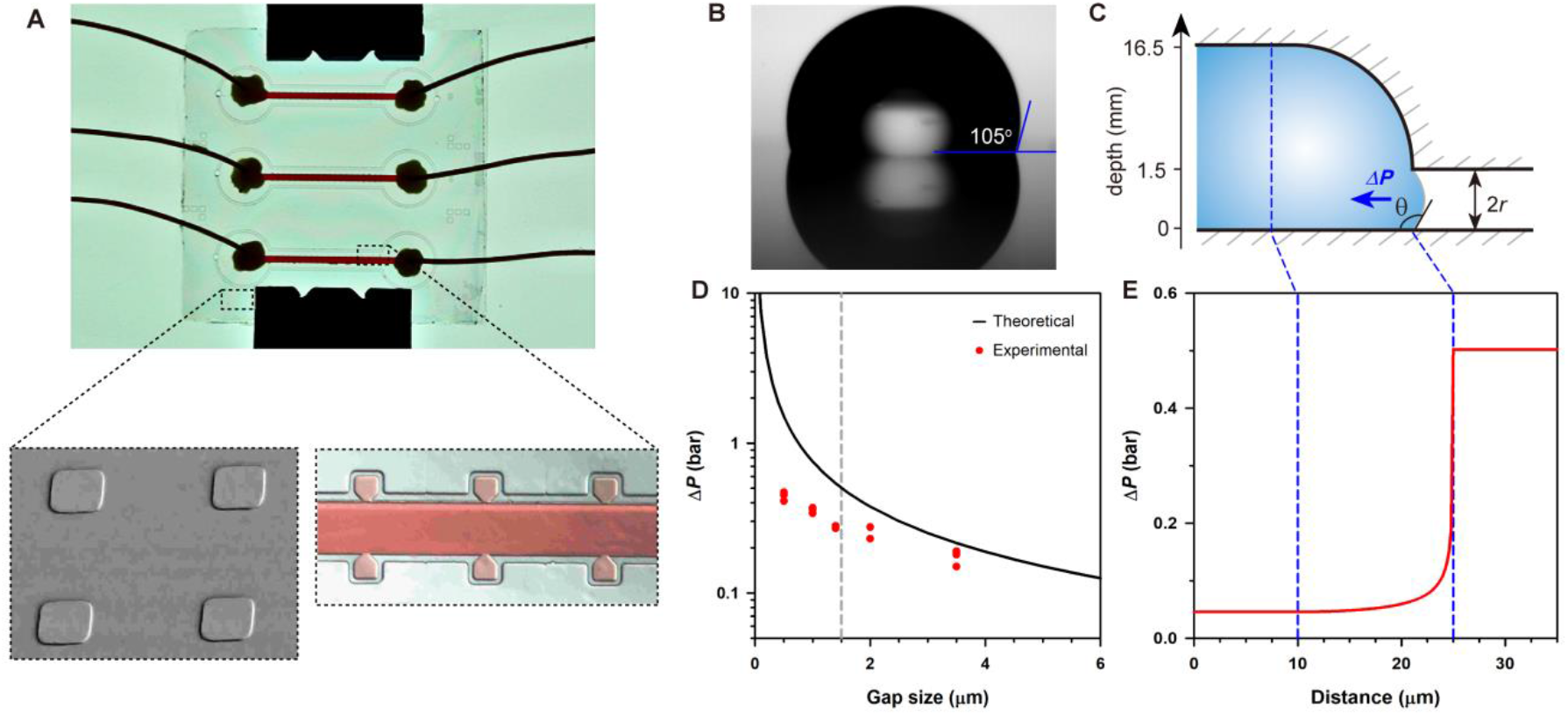
(A) A picture of the assembled device. 1.5-µm height nanogaps on the bottom plate were observed via scanning electron microscopy. An assembled device filled with red dye and zoom-in view of the channel with red dye. (B) The silanized glass plates of the device have a contact angle of 105° with deionized water. (C) Side view of the water-air interface between the microwell and the gas-filled chamber showing the direction of surface tension of solution at the edge of the microwells. (D) Relationship between surface tension and gap size at the water-air interface, and (E) the surface tension distribution along the microwell.

The interface was stable for continuous growth and observation without deleterious drift or shift.

We regard the liquid surface as a spherical surface so that the capillary pressure Δ*P* can be derived from the following equations:

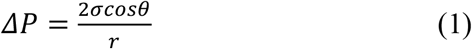

where, *σ* is the liquid surface tension (7.28 × 10^−2^ N/m); *θ* is the contact angle between the liquid surface and the solid plate; maximum value is 105°; the radius (*r*) of it equals to half of the gap height (0.75 µm). Thus, the capillary pressure Δ*P* can be calculated to be 5.03 × 10^4^ Pa, which is large enough to form a stable gas-liquid interface (Fig. 2D, E). *Streptomyces* spores were appropriately diluted and loaded into microwells to achieve single spore isolation in microwells following the Poisson distribution. Spores can germinate, branch to form vegetative hyphae in microwells, and later pass through the gap and differentiate into aerial hyphae in air-filled chambers that eventually develops into mature spores. The lifecycle of *Streptomyces* can last for several days, and thus we infused culture media continuously from the channel to guarantee adequate nutrient supply. The mycelia in the microwells would not be disturbed because of the narrow joint between the channel and the microwells. The entire developmental process could be monitored using an inverted microscope.

### Lifecycle observation of *Streptomyces coelicolor* on-chip culture in liquid medium

*S. coelicolor* is a model organism of *Streptomyces*, and the complete genome of the type strain *S. coelicolor* M145 has been sequenced; it is used in many studies of *Streptomyces* growth and development (2). We cultivated *S. coelicolor* in liquid minimal medium on the microfluidic device, and observed its entire lifespan (Fig. S1, Movie S1). After 9 h of dormancy, the spore emerged from one germ tube, which prolonged and formed branches. Each branch showed apical growth, indicating that the group of cells grew at an exponential phase in the microwell. The hyphae could spread randomly in liquid medium because there was no solid substrate confinement. The hyphae gradually approached the water-air interface, broke the surface tension, and grew into the air-filled chamber at 28 h (growth almost perpendicular to the edge of the microwell). The aerial hyphae progressively elongated and formed branches in all directions. There were curls and spirals at the end of the hyphae. Meanwhile, the vegetative hyphae developed many layers, and eventually almost filled the entire microwell. The vegetative hyphae and the aerial hyphae stopped growing after 60 hours (Movie S1).

When cultivated in the flask-scale liquid medium, aerial hyphae formation and sporulation are blocked in most *Streptomyces* strains (16), but when cultured in bioreactors, some strains may be able to sporulate due to stress conditions such as strong agitation (17). It has been suggested that nutrient depletion and the reuse of materials led to the hyphae differentiation in liquid medium (18), and programmed cell death also trigger the differentiation process in liquid and solid media (13). Although the specific signals are unclear, *N*-acetylglucosamine produced by the decomposition of peptidoglycan may be one of the signals (19). However, small-scale single-cell development has not been carefully illustrated, and differentiation might be limited. In this study, we cultivated *S. coelicolor* in a microfluidic device and found that vegetative hyphae did not lyse; rather, they continually grew even after the emergence of aerial hyphae. The culture media were supplied continuously in the microfluidic device such that the nutrition is not exhausted, indicating that the differentiation phenomenon may not be necessarily correlated with nutrient depletion.

### Differentiation of *S. coelicolor* on-chip culture in liquid YEME medium

*S. coelicolor* can form aerial hyphae and spores in standing liquid cultures with minimal media but not with complete media (20). Here, we inoculated single spores in microwells with nutrient-rich YEME medium and cultivated the samples several days to test whether they can differentiate (Fig. 3). The results showed that *S. coelicolor* still has a complete lifecycle in liquid YEME medium including vegetative hyphae in microwells (Fig. 3A), aerial hyphae outside the well (Fig. 3C), and mature spores (Fig. 3D). Scanning electron microscopy analysis revealed that the hyphae in the microwells had a relatively smooth surface (Fig. 3B). Hyphae in the chamber had a layer of well-organized hydrophobic proteins (21) (Fig. 3C) with compartments between each spore (Fig. 3D). These results are consistent with the development of *S. coelicolor* grown on solid plates and previous reports on the microscopic feature of hydrophobic proteins (22).

**FIG 3.**
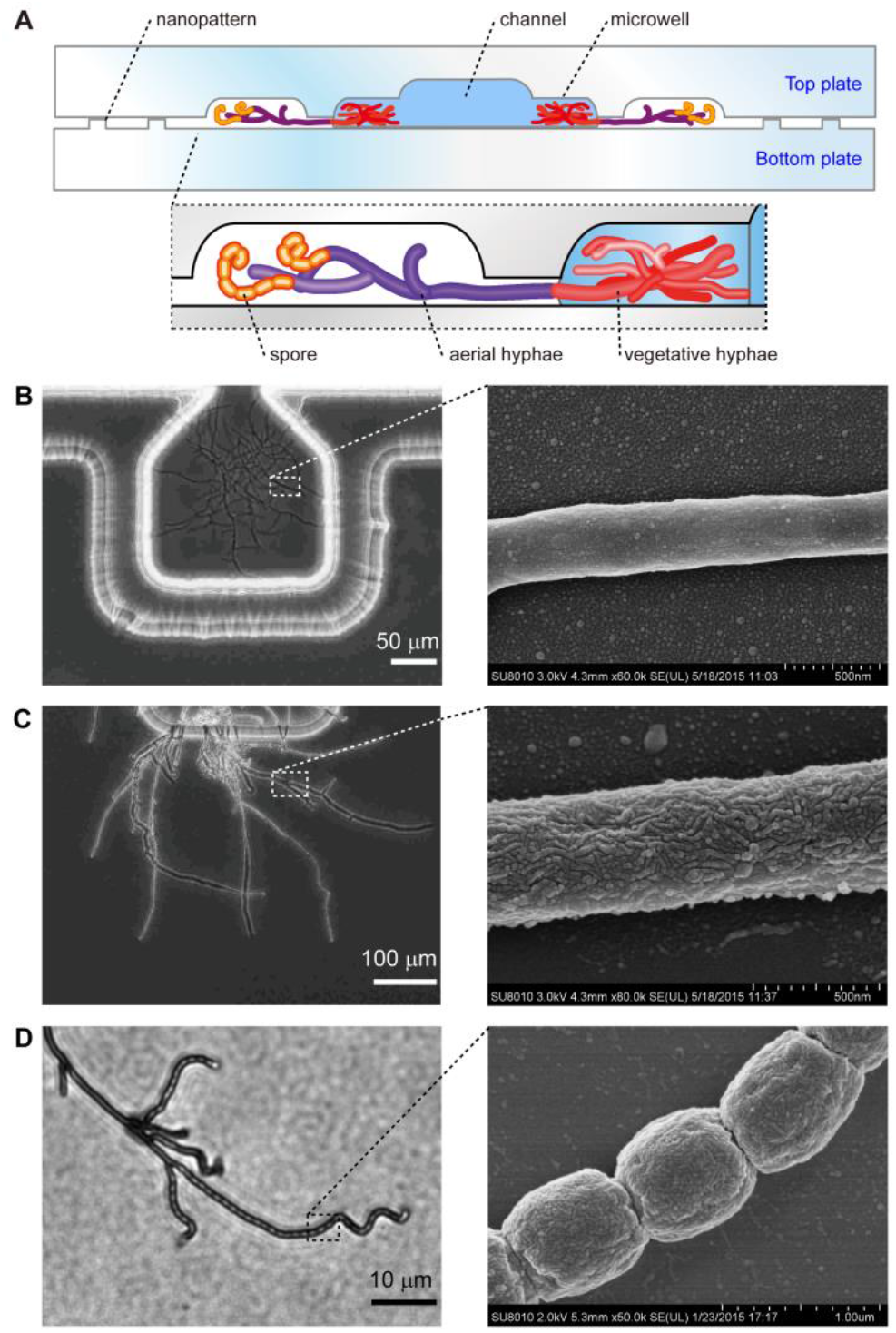
Development of *S. coelicolor* cultivated in a microfluidic device. (A) Schematic of sectional view of the device with *Streptomyces* development. (B-D) The vegetative hyphae (B), aerial hyphae (C), and spores (D) of *S. coelicolor* were observed by optical microscopy (left panel) and electron scanning microscopy (right panel), respectively.

Accordingly, *S. coelicolor* has entire lifecycles in the liquid environment regardless of the nutrient status. An earlier study showed that the expression of most genes is comparable between liquid and solid cultures, including genes involved in the hydrophobic cover formation and even a few genes regulating the early stages of sporulation (23). Genes involved in the final stages of hydrophobic cover/spore maturation are up-regulated in solid cultures compared with liquid cultures. These findings suggest that *S. coelicolor* can differentiate in both solid and liquid cultures. Transcripts and proteins are ready before aerial hyphae formation. Once *S. coelicolor* senses the existence of air, they begin to grow aerial hyphae and develop into mature spores. In standing liquid cultures, there may be a physical constraint that hinders the formation of aerial hyphae: The nutrient-rich media contains more complex ingredients, which is likely to attach to the hyphae surface and reduce the hydrophobicity of the hyphae, making it difficult for the aerial hyphae to erect.

Interestingly, we observed the merging of aerial hyphae when *S. coelicolor* was cultivated in the microfluidic chip (Fig. 4, Movie S2). This universal phenomenon in Streptomycetes is called hyphal anastomosis (or hyphal fusion), which was firstly confirmed in *S. scabies* (24), and is considered to be very important for intra-hyphal communication, nutrients and water translocation, and general homeostasis within a colony (25). There is hypha-to-hypha fusion, two hyphal tips grow towards each other until contact and fuse (Fig. 4A), as well as hypha-to-peg or hypha-to-side fusion, a hyphal tip approaches the side of another existing hypha and directly fuse with it (Fig. 4B, C) occurred in this study.

**FIG 4.**
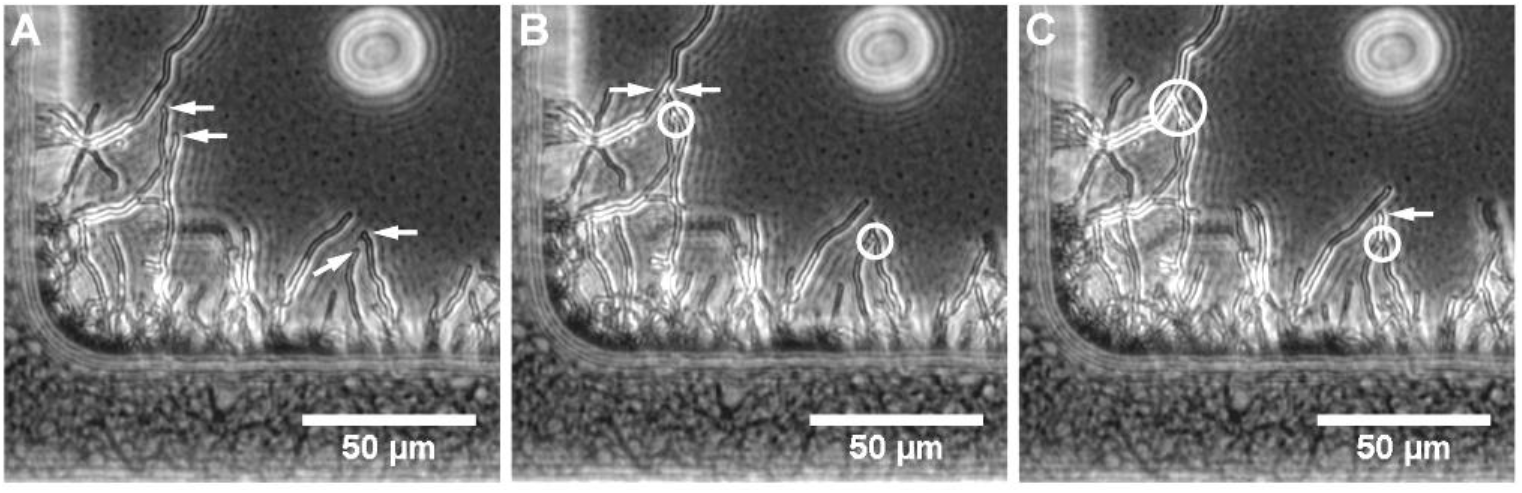
Hyphal anastomosis (fusion) in *S. coelicolor*. Some hyphal tips (arrowed) approaching for fusion (A). (B) shows fused hyphal branches (circled). A fused hyphal tip is growing towards a hyphal peg for subsequently fusion (C).

### Comparison of *S. griseus* wild type and mutant

To evaluate whether the differentiation in microfluidic liquid culture approach with *Streptomyces* can be generalized, another model organism *S. griseus* was cultivated in the microfluidic chip to observe its differentiation in liquid cultures. We cultured *S. griseus* in liquid MM medium and YEME medium, respectively, and observed its three lifespan stages through optical microscopy and electron microscopy (Fig. S2). The results confirmed that *S. griseus* can pass through its whole lifecycle in both liquid cultures.

The differentiation mechanism of *S. griseus* on the solid plate has been well-studied. The entire process begins with the expression of *afsA* that controls the synthesis of A-Factor, a type of *γ*-butyrolactones known as microbial hormones. A-Factor is essential in the regulatory pathways of sporulation (26), it can bind to A-Factor receptor protein ArpA and relieves the suppression of ArpA to *adpA*. AdpA can then stimulate a series of responses involving morphological development and secondary metabolism (Fig. S3A). Genes involved in the formation of aerial hyphae and spore, including *ssgA* (27), *adsA* (26), *amfR* (28), as well as extracellular proteases (29, 30) and protease inhibitor encoding genes (31) are regulated by AdpA. When *afsA* is knocked down, the mutant cannot form aerial hyphae on solid YEME plate. We constructed a *S. griseus* Δ*afsA* mutant via genetical engineering to compare the mechanism of differentiation between solid and liquid cultures (Fig. S3).

We inoculated the mutant on solid YEME medium and cultivated it for several days. Compared with wild-type, the mutant strain can neither develop aerial hyphae nor pigmented spores (Fig. 5B). When cultured on a microfluidic device, the hyphae were mainly in microwells with very few hyphae outside microwells. These were very short even after being cultivated for several days. They could not form spores (Fig. 5D).

**FIG 5.**
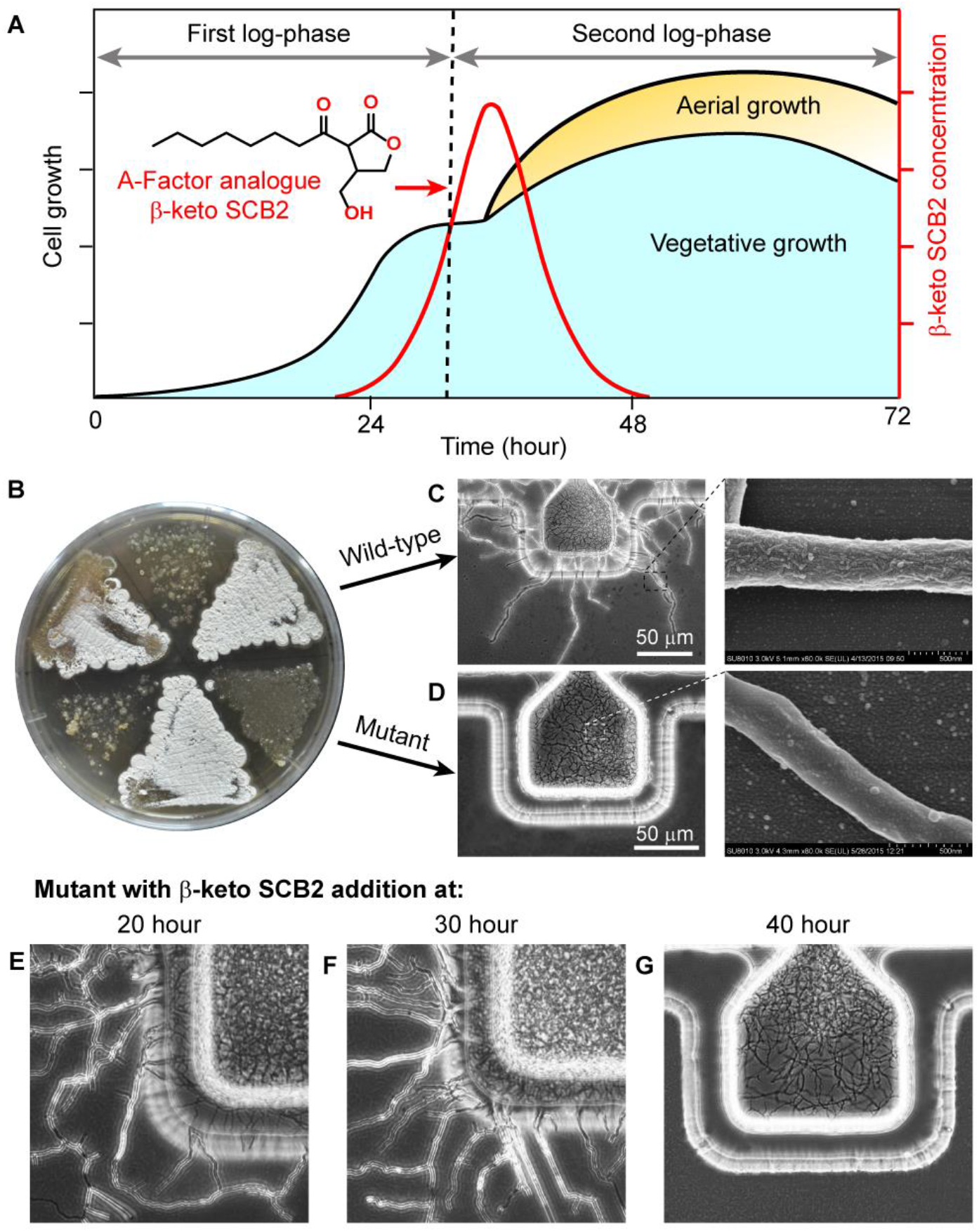
Biosynthesis of A-Factor in a growth-dependent manner (A) and its role in the development of *S. griseus* (B–E). Phenotypes of *S. griseus* wide-type and Δ*afsA* mutant on solid plate (B). Aerial hyphae of *S. griseus* wide-type (C) and Δ*afsA* mutant (D) cultivated in a microfluidic device were studied via optical microscopy and electron scanning microscopy. Feeding of A-Factor analogue and recovery of Δ*afsA* mutant phenotype at 20 h (E), 30 h (F), and 40 h (G) after cultivation.

Scanning electron microscopy showed that the hyphae surface of *S. griseus* Δ*afsA* mutant was relatively smooth (Fig. 5D). Thus, we inferred that these short hyphae are still vegetative hyphae. While they emerged out of the microwell, they could not develop further. The phenotype of *S. griseus* Δ*afsA* mutant grown in liquid culture was similar to that grown on the solid plate. These results demonstrate the consistency in observation of differentiation behavior of *S. griseus* wild type and mutant strains between microfluidic and bulk cultures.

### Feeding of A-Factor analogue and recovery of the mutant phenotype

The A-Factor analogue *β*-keto SCB2 was chemically synthesized as described above and added to the culture medium to examine whether the mutant could recover a wild-type phenotype. Previous studies showed that A-Factor is chemically unstable with a half-life of several hours. The production of A-Factor is growth-dependent and accumulated with the growth of hyphae (Fig. 5A). It reaches peak concentration of 25–30 ng/mL and rapidly decreases thereafter.(15) According to the development of *S. griseus* and the instability of A-Factor, we fed *β*-keto SCB2 to the Δ*afsA* mutant at several time points in the lifecycle. When *β*-keto SCB2 was added at 20 h and 30 h after inoculation, the mutant could form aerial hyphae and spores (Fig. 5E, F).

Scanning electron microscopy further confirmed that there were hydrophobic proteins on the surfaces of hyphae and spores observed from *S. griseus* Δ*afsA* mutant on device with *β*-keto SCB2 supplied at 30 h after initial cultivation. (Fig. S4). When *β*-keto SCB2 was added at 40 h after inoculation, the mutant could no longer form aerial hyphae (Fig. 5G). These results are consistent with previous studies showing that timing is critical for A-Factor’s switching function (Fig. 5A) (15). There is a decision phase at the middle of the exponential growth, which is an A-Factor-sensitive period. The exogenous addition of A-Factor after this time can no longer influence morphological differentiation (15). We conclude that the microfluidic chip is compatible with A-Factor assay by introducing the test compound at defined time point, and demonstrated the timing-sensitive effect of A-Factor on morphological differentiation of mutant deficient in A-Factor synthesis, which is in good agreement with the classical literature reports.

### Mechanism of water-air interface differentiation and possible applications of the microfluidic device

To confirm whether the differentiation of *Streptomyces* is different between that grown on solid plates and that grown in liquid media, we conducted an A-Factor-regulated experiment in *S. griseus* using the microfluidic-based device. Timing studies on the microfluidic device and scanning electron microscopy showed that the differentiation mechanism of *S. griseus* in the liquid environment is the same as that on solid plates. We also found that sporulation is not correlated with lysis of vegetative hyphae suggesting that genes encoding extracellular proteases and protease inhibitors may not be necessary for differentiation.

Differentiation mechanisms of *Streptomyces* are essential for the study of secondary metabolism. Using this microfluidic device, we successfully restored the wild-type phenotype of *S. griseus* Δ*afsA* mutant by addition of the A-Factor analogue *β*-keto SCB2. Mechanisms by which other chemical molecules affect the morphological differentiation and secondary metabolism of *Streptomyces* can also be studied through this device with significantly reduced consumption of compound of interest by virtue of miniaturization. Previous studies on the differentiation of *Streptomyces* in liquid media mainly focused on the analysis of pellets and clumps formation (32). Some strains require the formation of pellets to produce secondary metabolites, such as *S. coelicolor* (undecylprodigiosin and actinorhodin) (13) and *S. olidensis* (retamycin) (33) while pellets and clumps formation reduce the antibiotic productions in *S. noursei* (nystatin) (34) and *S. fradiae* (tylosin) (35). The microfluidic device we developed will help to establish the developmental model of different *Streptomyces* strains in liquid cultures, which will be beneficial to the optimization of industrial fermentation.

In addition, a previous study found that *S. coelicolor* was able to produce several secondary metabolites during its germination: albaflavenone (antibacterial activity against *Bacillus subtilis*), the polyketide germicidin A, and chalcone (inhibits germination) (36). As the whole lifecycle of *Streptomyces* can be observed by the microfluidic device, functions of secondary metabolites produced by *Streptomyces* strains in their early stages of growth can also be further studied.

## CONCLUSION

We developed a microfluidic device for cultivation and the entire lifecycle observation of *Streptomyces*, a dominant paradigm in bacterial multicellularity evolution. Compared with traditional methods, this microfluidic device can achieve single-cell long-term dynamic cultivation and observation enabled by nanogap-stabilized air-water interfaces. Two model strains of *Streptomyces* (*S. coelicolor* and *S. griseus*) were cultivated in the microfluidic device at single-cell/spore resolution. Cellular development and differentiation of the entire *Streptomyces* lifecycle was monitored with microscopy and further characterized with scanning electron microscopy on the microfluidic device for the first time. We also studied *Streptomyces*’ development under different nutrient conditions or chemical stimuli, and we may also use it to investigate the cell-cell interaction between *Streptomyces* and pathogenic bacteria.

We found that aerial mycelia may grow at a very early stage, and such formation may not be necessarily correlated with nutrient depletion. Moreover, we constructed a *S. griseus* Δ*afsA* mutant (exogenously supplemented A-Factor analog to the mutant) and compared the differentiation mechanism between solid and microscale liquid environments. We inferred that *Streptomyces* still has the entire lifecycle in a microscale liquid environment, and the differentiation mechanism is the same as that on solid plates. These new observations shed light on the understanding of *Streptomyces*’ multicellular development and differentiation. Although the current biological data are findings and conclusion derived from visual inspections of images, other scientific analysis and representation also can be achieved by further image analysis, for example, growth of the different morphological states (Fig. 3) could be characterized and compared by derived growth rates from image data of filamenting organisms (11).

Overall, we anticipate that our new method provides a better platform for the study of *Streptomyces*’ development in the natural porous and moist soil environment. The microwell arrays with stabilized air-water interfaces can mimic ecological niches and help us identify single-cell heterogeneity. *Streptomyces*’ complete lifecycle on the microfluidic device may also awaken cryptic secondary metabolite gene clusters for the secretion of secondary metabolites and lead to the discovery of novel antibiotics for combating global crisis of antimicrobial resistance.

## MATERIALS AND METHODS

### Bacterial strains and materials

The microbial strains used in this work include *Streptomyces coelicolor* M145, *Streptomyces griseus* IFO 13350, and *S. griseus* Δ*afsA* mutant. These strains were cultured on the Mannitol-Soy agar plate at 28 °C for about a week to allow spore germination. The spores were harvested by sterile cotton swabs and suspended in the sterilized culture medium. The suspension was filtered through a filter tube filled with cotton wool to remove aerial hyphae. The OD_600_ of the spore suspension was adjusted to 0.15 to ensure that most of the microwells had a single spore. Liquid minimum (MM) medium and yeast extract-malt extract (YEME) medium were used for on-chip cultivation.

### Fabrication of the device

The microfluidic device was made of two glass plates and fabricated by standard photolithography as well as wet chemical etching techniques (37). The photomasks were designed using AutoCAD (San Rafael, CA) and ordered from MicroCAD photomask Co. Ltd. (Shenzhen, China). The top plate has a 55-μm-deep channel, with 40 microwells symmetrically distributed along the channel with a volume of 0.27 nL individually. The bottom plate consists of an array of nanogaps of 1.5 μm high (Fig. 1A and Fig. 2A). The top plate has two access holes drilled by a diamond drill bit 0.8 mm in diameter. The glass plates were cleaned with ethanol, oxidized in a plasma cleaner, and silanized by 1*H*,1*H*,2*H*,2*H*-perfluorooctyl trichlorosilane.

### Device operation and cell cultivation

The glass chip was thoroughly cleaned with ethanol and tightly clamped by clips. The spore suspension was aspirated into a pipette and loaded into the channel leading to the microwells (Fig. 1B). Then suspension in the channel was aspirated from the outlet to remove excess spores to prevent channel block caused by hyphae growth, but medium and spores in microwells could be retained (Fig. 1B). Two syringes were connected to the device by Teflon tubing to infuse culture medium continuously for long-term cultivation (Fig. 1B). The device was placed under an inverted microscope to capture pictures every hour. A CO_2_ microscope cage incubator was placed around the microscope to maintain the temperature at 28 °C for *Streptomyces* cultivation.

### Scanning Electron Microscopy

When cultivation terminated, the device was transferred to a freezer at -20 °C for one minute to freeze the sample, so that the hyphae could not move when we opened the device. The device was disassembled quickly, and the top plate was cut into 0.5 × 0.7 cm pieces and fixed in 3% glutaraldehyde overnight at 4 °C to maintain the bacteria’s physiological status. The sample was then washed with DI water to remove glutaraldehyde, dehydrated in an ethanol series (50%, 70%, 85%, 95%, and 100%), critical point dried, and sputter-coated with platinum under vacuum. The sample was observed under a scanning electron microscope.

### Synthesis of A-Factor analogue *β*-keto SCB2

*S. coelicolor* butanolides (SCBs) are *γ*-butyrolactones from *S. coelicolor*, and *β*-keto-SCB2 is a stereoisomer of A-Factor (38, 39). The synthesis of *β*-keto-SCB2 was began with methyl 3-oxocyclobutane-1-carboxylate to methyl 5-oxotetrahydrofuran-3-carboxylate by the Baeyer-Villiger oxidation (40-43). Methyl 5-oxotetrahydrofuran-3-carboxylate was then reduced to 4-(hydroxymethyl)-dihydrofuran-2(3*H*)-one by the addition of NaBH_4_ followed by protection of hydroxyl group with *tert*-butyldimethylsilyl (TBS) chloride. Octanoyl chloride was slowly added to react with 4-(((*tert*-butyldimethylsilyl)oxy)methyl)dihydrofuran-2(3*H*)-one to give 4-(((*tert*-butyldimethylsilyl)oxy)methyl)-3-octanoyldihydrofuran-2-(3*H*)-one. The silyl protecting group was then removed with tetrabutylammonium fluoride to afford *β*-keto-SCB2. Mass spectrometric analysis characterized the synthetic products. See SI for detailed synthetic steps.

## SUPPLEMENTAL MATERIAL

Supplemental material contains:

Supplementary figures on lifecycle of *S. griseus* (FIG S1-S4), and the synthesis

process of A-Factor analogue *β*-keto SCB2, PDF file, 1.54 MB.

Movie S1 shows lifecycle of *S. coelicolor*, AVI file, 3.81 MB.

Movie S2 shows hyphal anastomosis of *S. coelicolor*, AVI file, 0.99 MB.

## ACKNOWLEDGMENTS

This work was supported by the National Natural Science Foundation of China (Nos. 31970091, 21822408 and 91951103), the program of China Ocean Mineral Resources R&D Association (No. DY135-B-02), the National High Technology Research and Development Program of China (No. 2018YFC0310703), and Senior User Project of RV KEXUE from Center for Ocean Mega-Science, Chinese Academy of Sciences (KEXUE2019GZ05).

## Table of Content Graphic

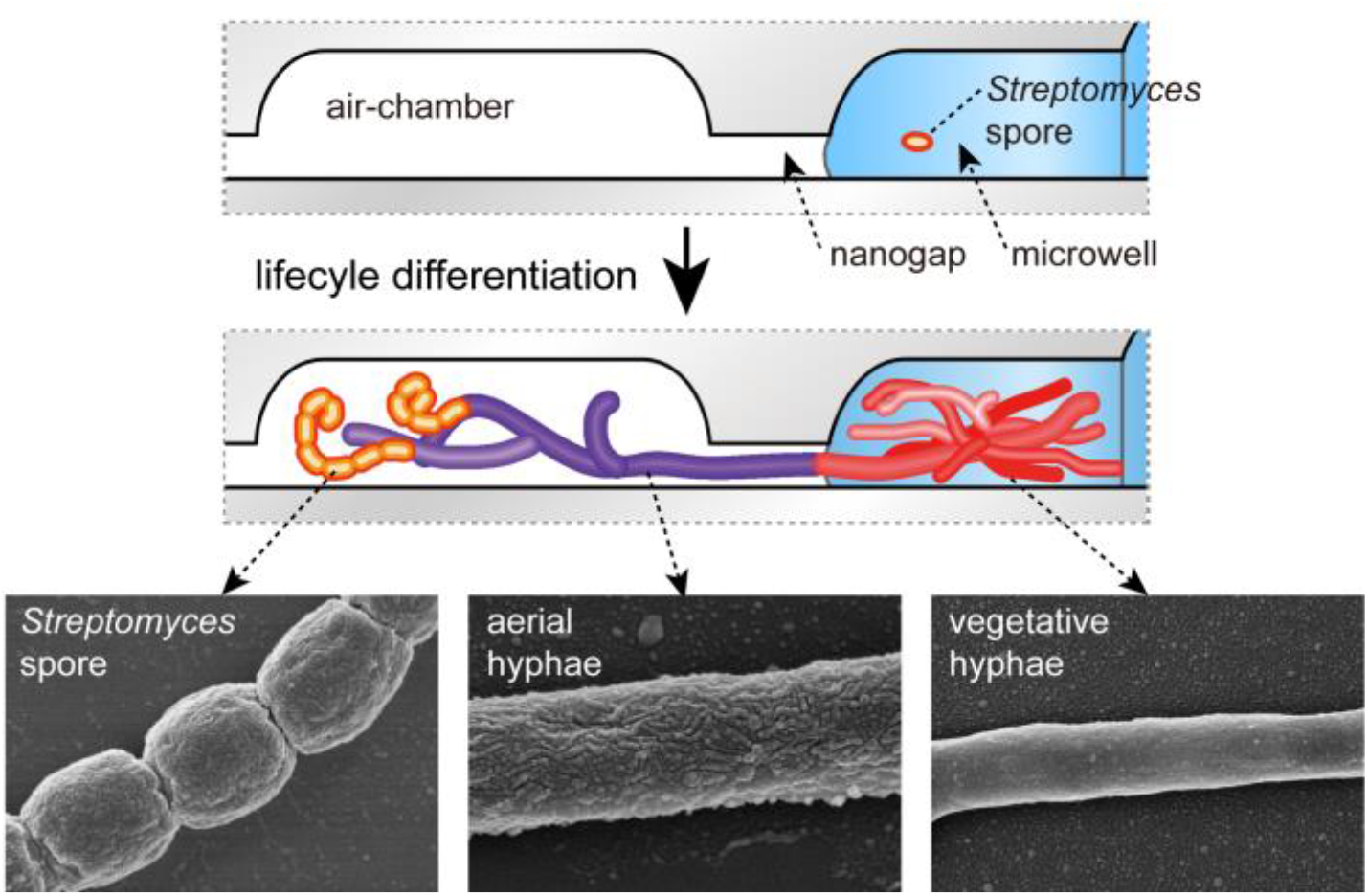

