## Supplementary material for "Microfluidic *Streptomyces* Cultivation for Whole Lifecycle Characterization and Phenotypic Assays Enabled by Nanogap-stabilized Air-Water Interface": Figure S1-4

**Construction of the *S. griseus* Δ*afsA* mutant.** The *idgS*-based blue-white screening system was used to construct the *S. griseus* Δ*afsA* mutant (1). To eliminate the polar effects, we disrupted the *afsA* gene along with the following gene *bprA* that was involved in the A-Factor analogue biosynthesis (Fig. 5A). Two 1.3-kb homologous arms flanking the *afsA* and *bprA* gene were obtained with the *S. griseus* IFO13350 genomic DNA as a template with the following primers afsA-LU (5'-CTTCCATGGGCACG C**CCTAGG**GCGCCTTCACCCGCTG ACT-3' with a *BlnI* site shown by bold letters), afsA-LD (5'-AGCGGTATCCAGGGGCCTA GGGCACCATCTCGATCCCCAC-3'), afsA-RU (5'-GGGACTCTGG GGTTCCAATTGT GGTGCCCACGGTCCAAGA-3' with a *MunI* site shown by underlining), and afsA-RD (5'-CTT GCTAGCAGATGTCAATTGGCCGAGGATGTTTACCCAGGTC-3'). The fragments were then cloned to the *MunI* and *BlnI* sites of pCIMt004 through a ligation-independent cloning strategy, resulting in pCIMt004_*afsA_bprA* (Fig. S3B). The construction of pCIMt004_*afsA_bprA* was verified by nucleotide sequencing. Plasmid pCIMt004_*afsA_bprA* was then introduced into *S. griseus* IFO 13350 by *E. coli-Streptomyces* conjugation. After blue-white screening, the apramycin resistant double-crossover mutants were picked up and confirmed by PCR with an *afsA*-LU and *afsA*-RD primer pair (Fig. S3C).

**Synthesis of A-Factor analogue *β*-keto SCB2 (2-7)**

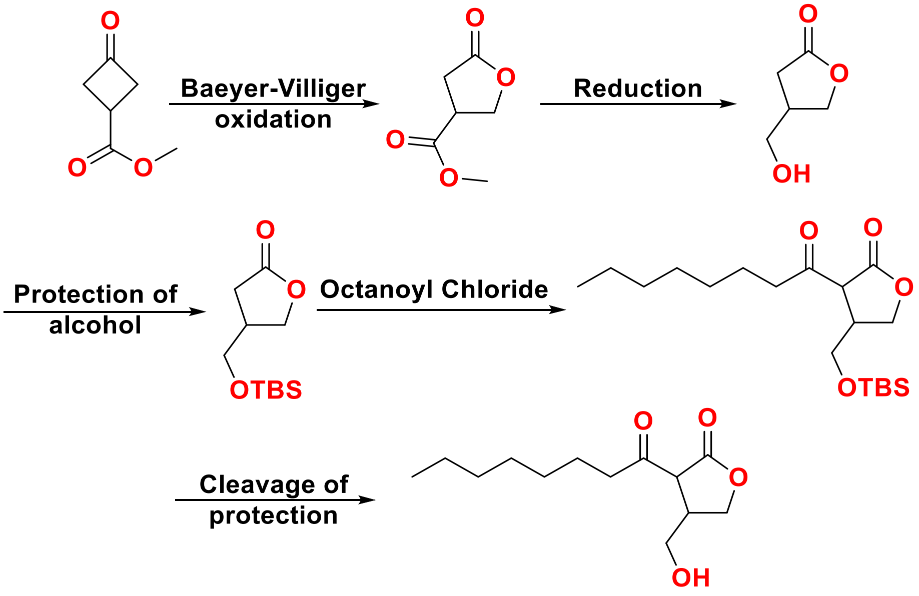


**(I) Methyl 5-oxotetrahydrofuran-3-carboxylate**

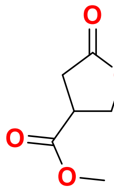

A dichloromethane (3 mL) solution of methyl 3-oxocyclobutane-1-carboxylate (0.1 g) at room temperature was added to $H_2O_2$ (30%, 0.25 g) with a few drops of acetic acid. The reaction mixture was then warmed to 50 °C overnight. The reaction mixture was quenched with saturated aqueous NaCl, extracted with dichloromethane, dried with $Na_2SO_4$, and concentrated under reduced pressure. The crude product was purified by silica gel chromatography to give methyl 5-oxotetrahydrofuran-3-carboxylate.

**(II) 4-(Hydroxymethyl)-dihydrofuran-2(3*H*)-one**

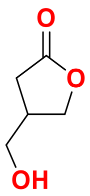


A THF (1 mL) solution of methyl 5-oxotetrahydrofuran-3-carboxylate (0.1 g) at 0 °C was added to $NaBH_4$. A small volume of MeOH was added dropwise, and the reaction mixture was warmed to room temperature for 15 min. The reaction mixture was quenched with saturated aqueous $NH_4Cl$, extracted with dichloromethane, dried with $Na_2SO_4$, and concentrated under reduced pressure. The crude product was purified by silica gel chromatography to give 4-(hydroxymethyl)- dihydrofuran-2(3*H*)-one.

**(III) 4-(((*tert*-Butyldimethylsilyl)oxy)methyl)dihydrofuran-2(3*H*)-one**

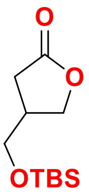


To a solution of 4-(hydroxymethyl)-dihydrofuran-2(3*H*)-one (20 mg) in 3 mL dichloromethane, TBSCl (38.8 mg), $Et_3N$(100 mg), and catalytic amounts of DMAP were added. After stirring for 36 h, the reaction mixture was quenched with saturated aqueous NaCl, extracted with dichloromethane, dried with $Na_2SO_4$, and concentrated under reduced pressure. The crude product was purified by silica gel chromatography to give 4-(((*tert*-butyldimethylsilyl)oxy)methyl)dihydrofuran-2(3*H*)-one.

**(IV) 4-(((*tert*-Butyldimethylsilyl)oxy)methyl)-3-octanoyldihydrofuran-2-(3*H*)-one**

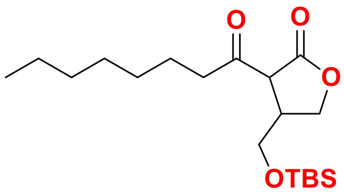


A solution of 4-(((*tert*-butyldimethylsilyl)oxy)methyl)dihydrofuran-2(3*H*)-one (40 mg) in THF (2 mL) was stirred at -78 °C for 5 min. LiHMDS (1 M in THF 0.3 mL) was then added dropwise to the reaction mixture and stirred for 30 min; octanoyl chloride (40 mg) was slowly added to the mixture, and then warmed to room temperature overnight. The reaction mixture was quenched with saturated aqueous NaCl, extracted with dichloromethane, dried with $Na_2SO_4$, and concentrated under reduced pressure. The crude product was purified by silica gel chromatography to give 4-(((*tert*-butyldimethylsilyl)oxy)methyl)-3-octanoyldihydrofuran-2-(3*H*)-one.

**(V) A-Factor analogue, β-keto SCB2**

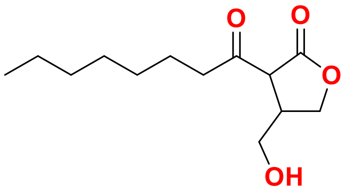


To a solution of 4-(((*tert*-butyldimethylsilyl)oxy)methyl)-3-octanoyldihydrofuran-2-(3*H*)-one in THF and acetic acid (1:1), TBAF (1 M in THF) solution was then added to the reaction mixture under ice-bath condition and then warmed to room temperature until the reaction was completed. The reaction mixture was quenched with saturated aqueous $NH_4Cl$, extracted with dichloromethane, dried with $Na_2SO_4$, and concentrated under reduced pressure. The crude product was purified using silica gel chromatography to give *β*-keto SCB2 (2-[1′-hydroxyoctyl]-3-hydroxymethylbutanolide).

HR-ESI-MS spectrum of synthetic products are shown as below:

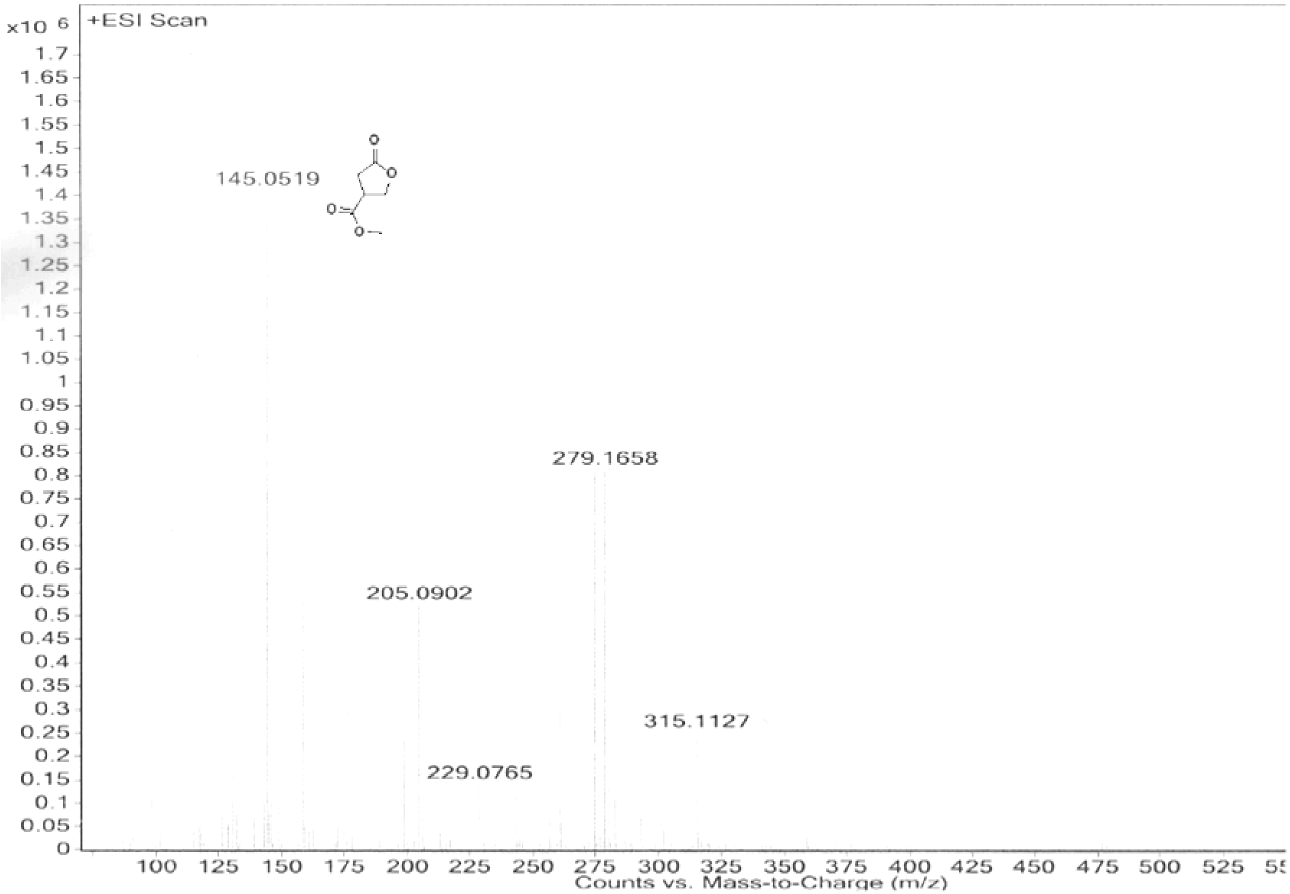

| | | | | | | | |
|---|---|---|---|---|---|---|---|
| **Sample Name** | GBL | **Position** | Vial 58 | **Instrument Name** | Instrument 1 | **User Name** | |
| **Inj Vol** | 60 | **InjPosition** | | **SampleType** | Sample | **IRM Calibration Status** | Success |
| **Data Filename** | GBL.d | **ACQ Method** | wwz-MeOH-LONGGRADE(+ | **Comment** | | **Acquired Time** | 8/28/2013 11:16:02 PM |

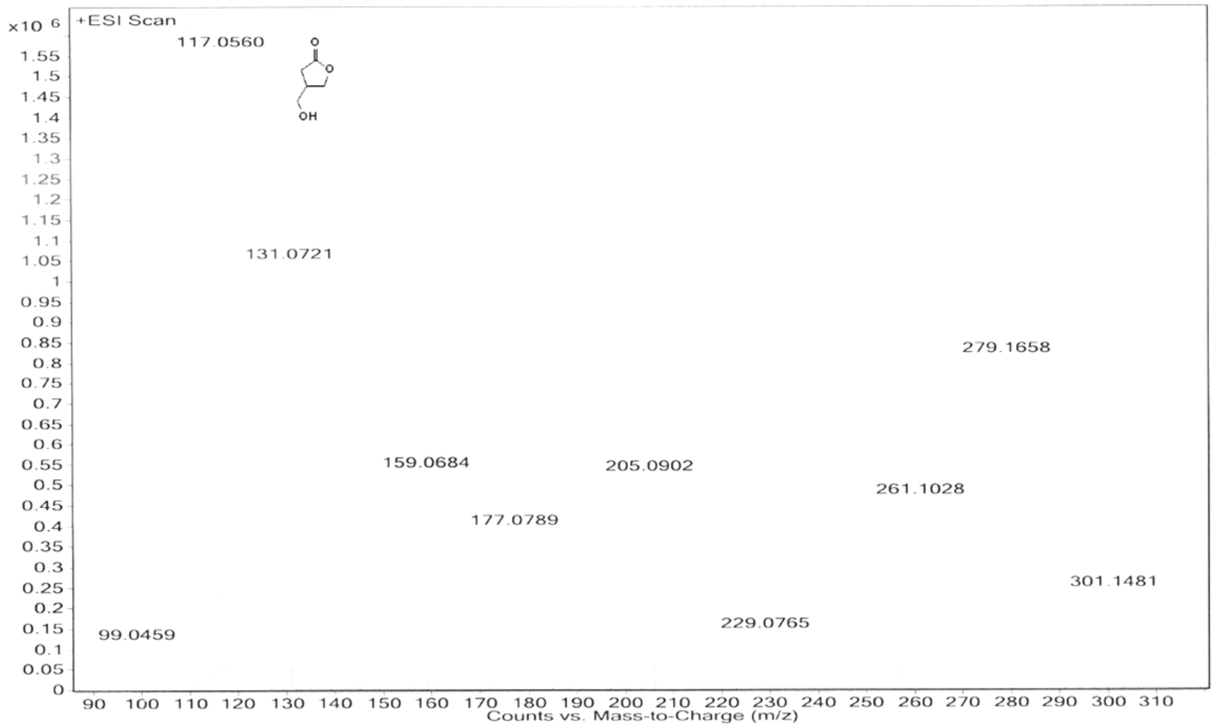


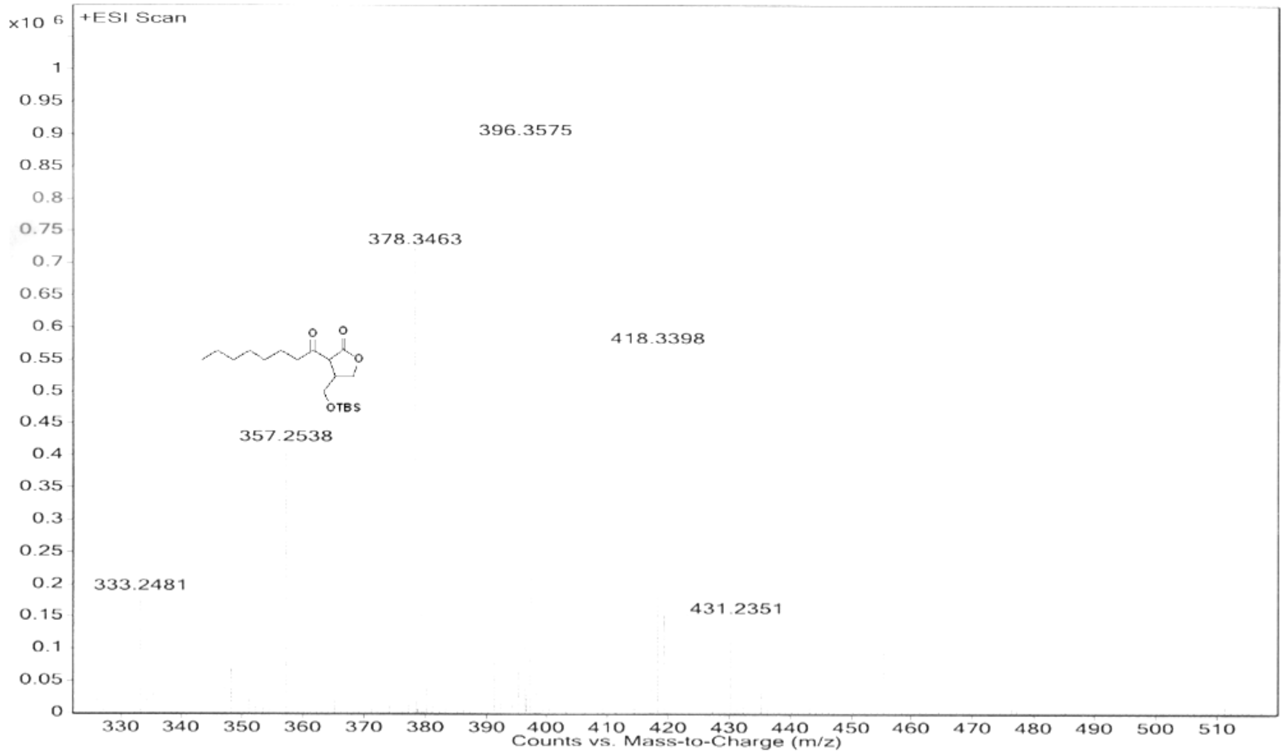

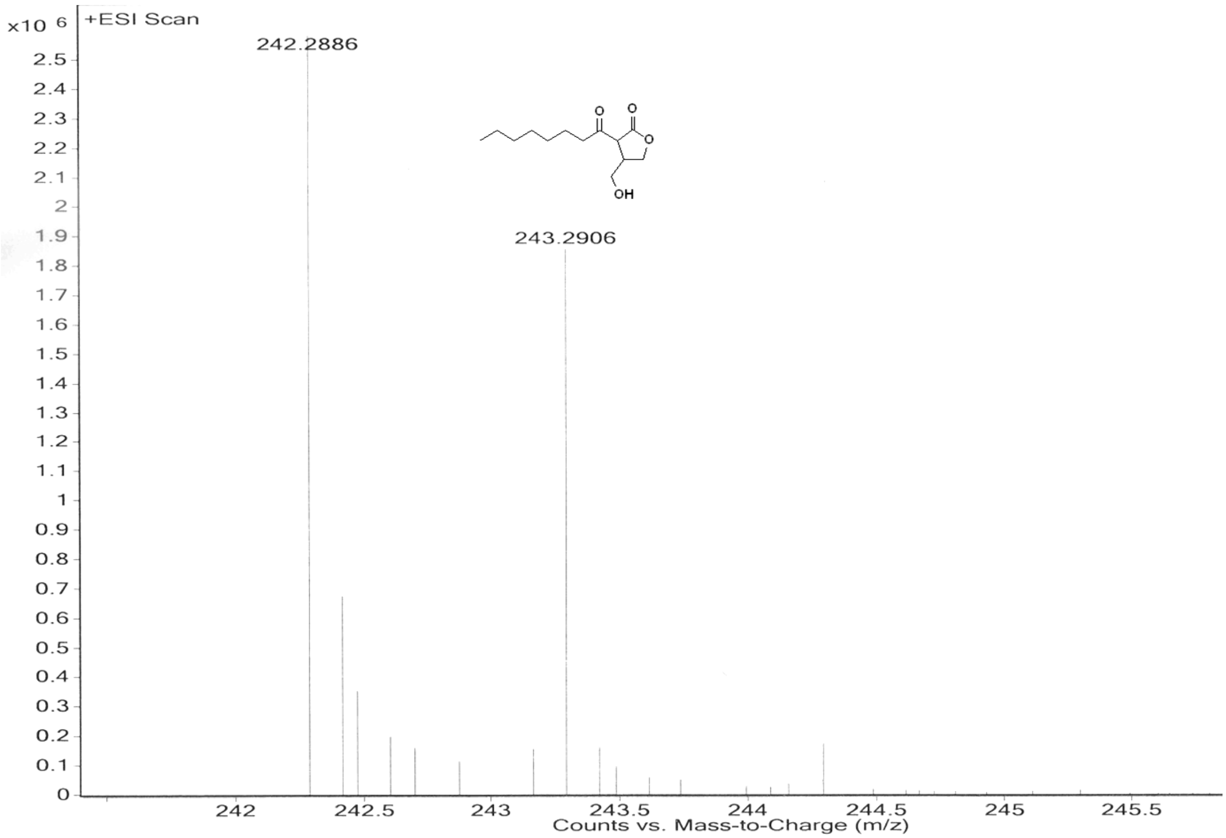

x10 6
+ESI Scan
242.2886
243.2906
OH
Counts vs. Mass-to-Charge (m/z)

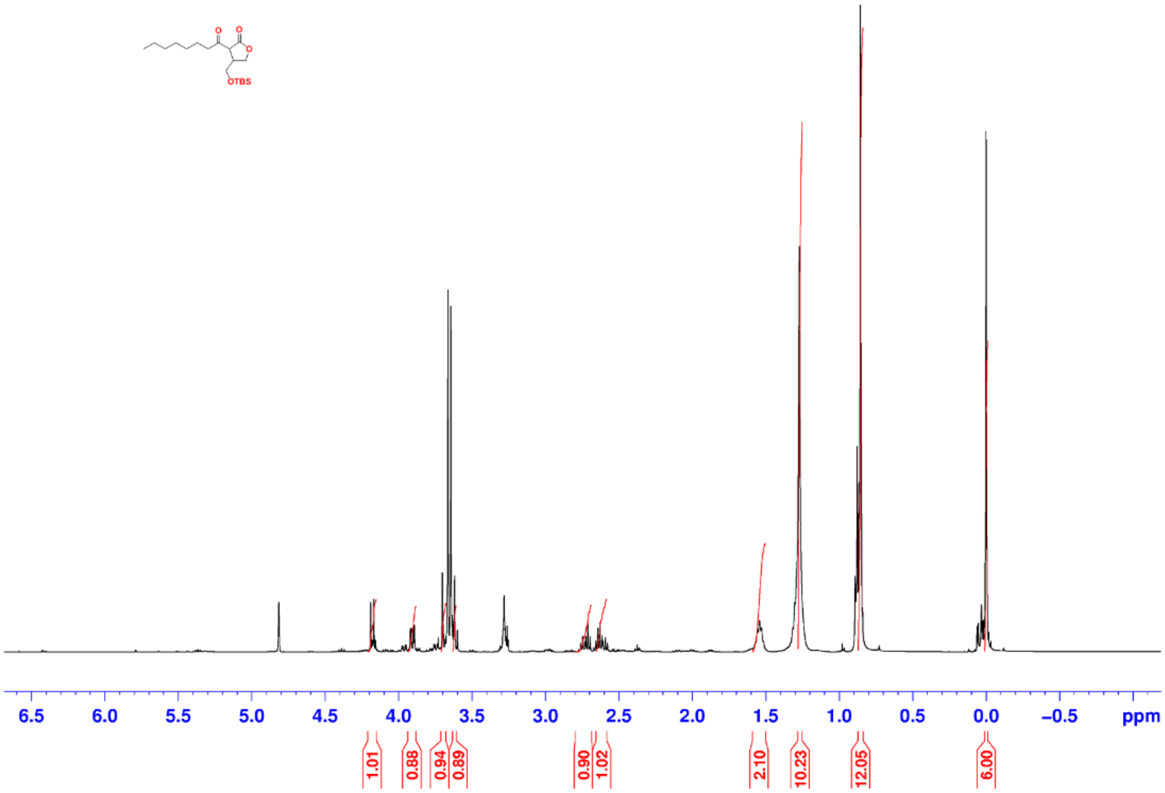

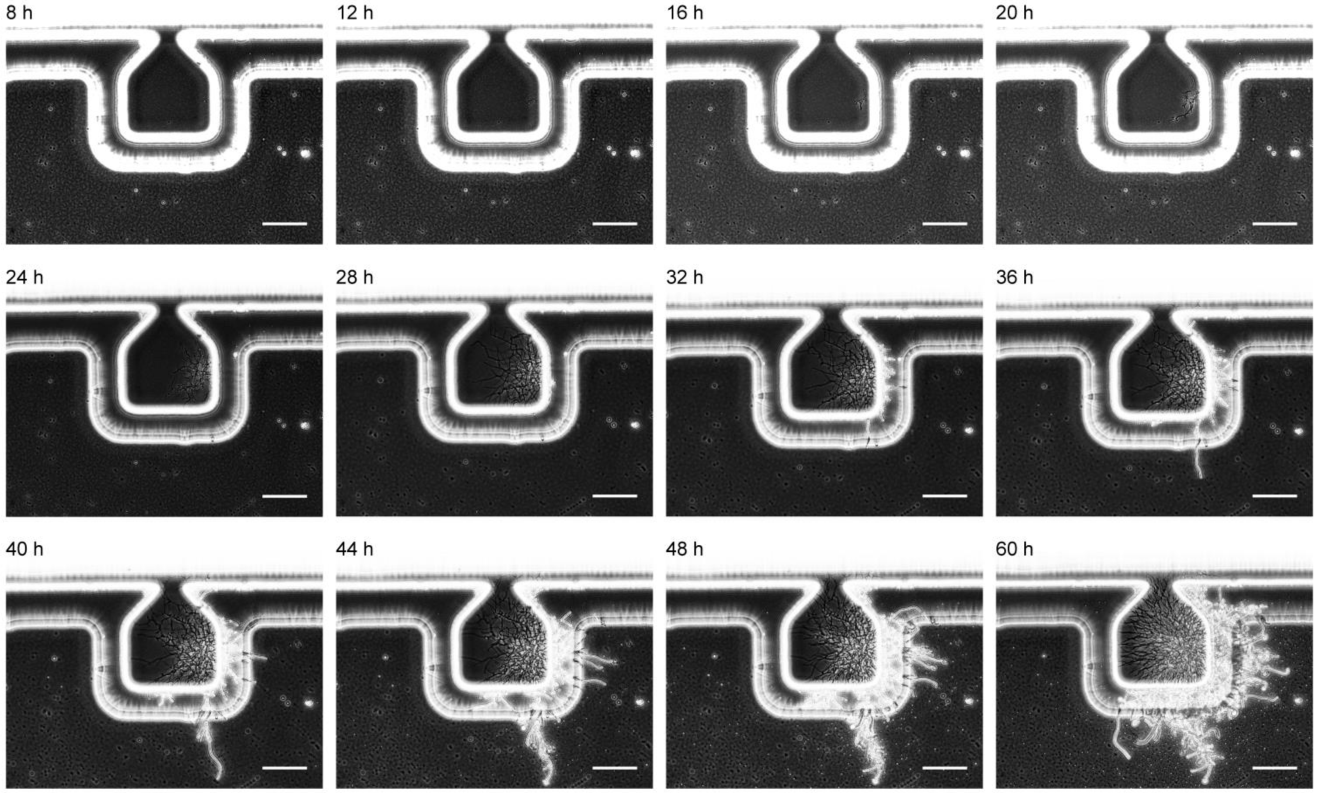


**FIG S1** Lifespan observation of *S. coelicolor*.

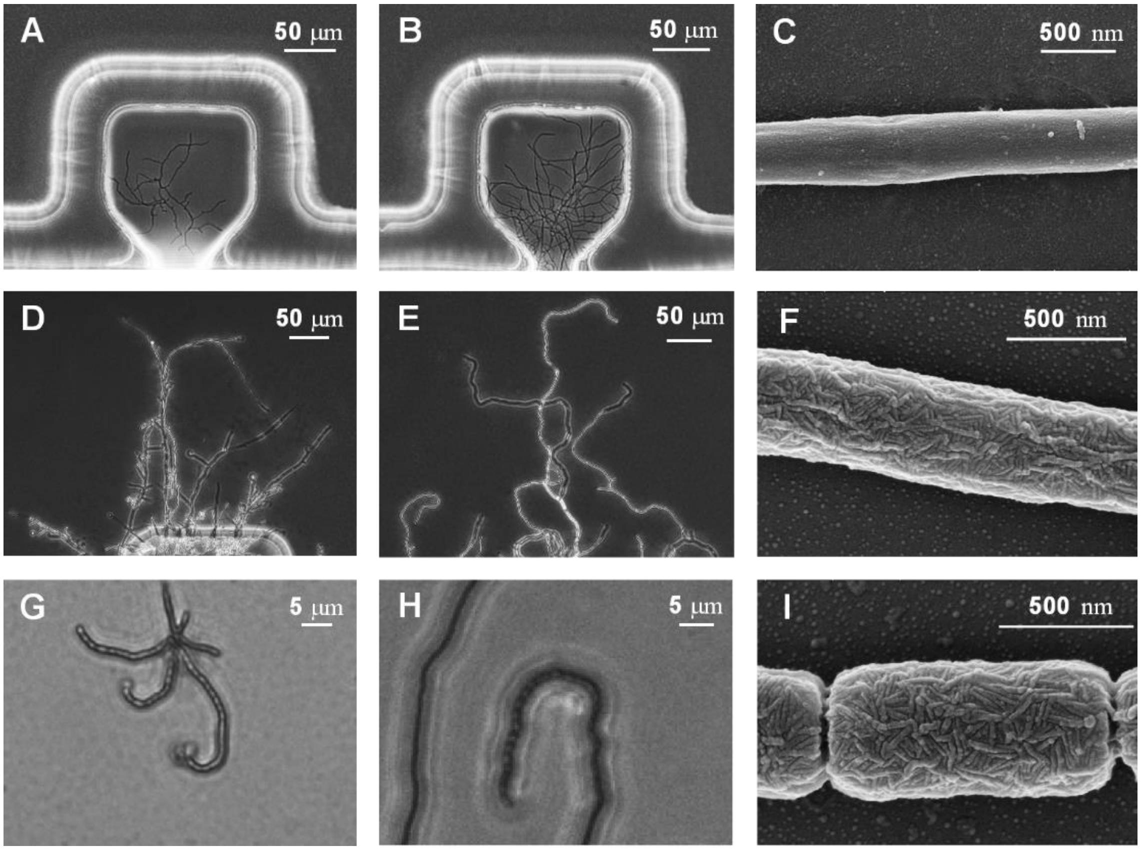


**FIG S2** Images of development of *Streptomyces griseus*. (A) Optical microscopy of vegetative hyphae cultivated with liquid MM media (B) Optical microscopy of vegetative hyphae cultivated with liquid YEME media. (C) Electron microscopy of vegetative hyphae. (D) Optical microscopy of aerial hyphae cultivated with liquid MM media. (E) Optical microscopy of aerial hyphae cultivated with liquid YEME media. (F) Electron microscopy of aerial hyphae. (G) Optical microscopy of spore cultivated with liquid MM media. (H) Optical microscopy of spores cultivated with liquid YEME media. (I) Electron microscopy of spores.

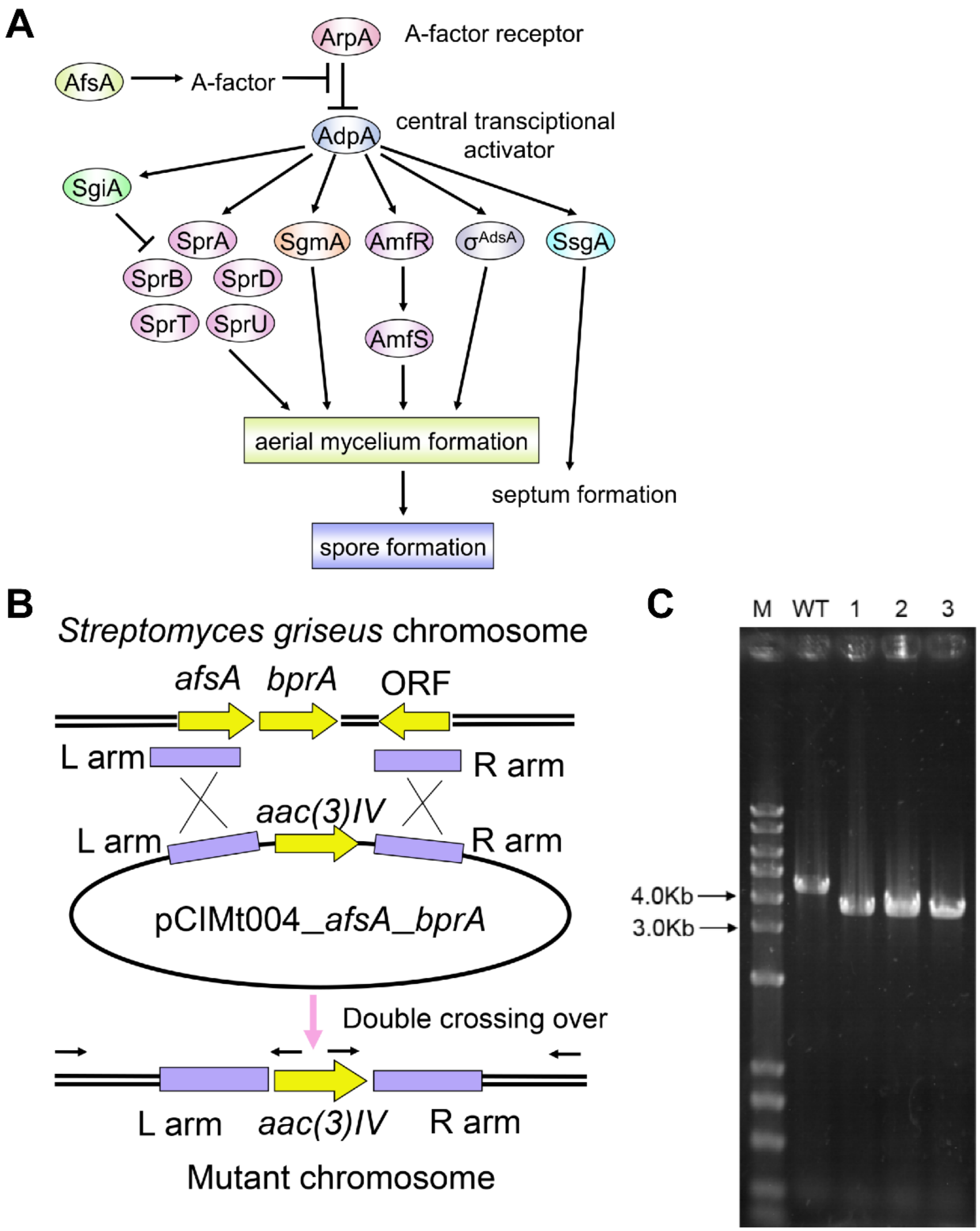


**FIG S3** (A) Illustration of A-Factor regulatory pathway leading to sporulation. (B) Illustration of *S. griseus* Δ*afsA* mutant construction; (C) electrophoresis of PCR products of *S. griseus* wide type and Δ*afsA* mutants.

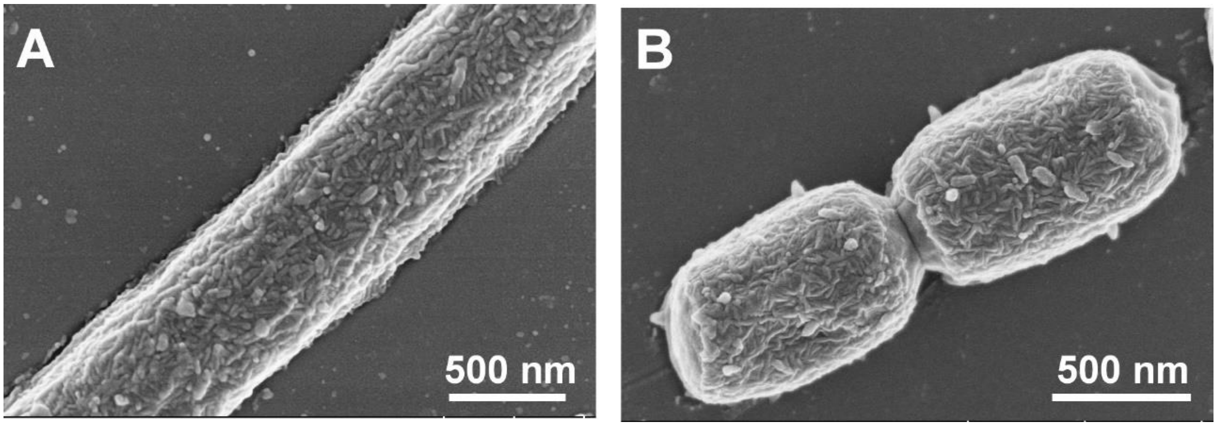


**FIG S4** Aerial hyphae and spores observed from *Streptomyces griseus* Δ*afsA* mutant on device with A-Factor analogue supplied at 30 hours after initial cultivation. Representative electron microscopy of aerial hyphae (A) and spores (B).
